# Decreased grey matter volume in mTBI patients with post-traumatic headache compared to headache-free mTBI patients and healthy controls: a longitudinal MRI study

**DOI:** 10.1101/487538

**Authors:** Shana A.B. Burrowes, Chandler Sours Rhodes, Timothy J. Meeker, Joel D. Greenspan, Rao P. Gullapalli, David A. Seminowicz

**Author notes:** Please address correspondences to: David A. Seminowicz, PhD, Department of Neural & Pain Sciences, University of Maryland School of Dentistry, 650 W. Baltimore Street, 8 South Baltimore, MD 21201.

## Abstract

**Background:** Traumatic brain injury (TBI) occurs in 1.7 million people annually and many patients go on to develop persistent disorders including post-traumatic headache (PTH). PTH is considered chronic if it continues past 3 months. In this study we aimed to identify changes in cerebral grey matter volume (GMV) associated with PTH in mild TBI patients.

**Methods:** 50 mTBI patients (31 Non-PTH; 19 PTH) underwent MRI scans: within 10 days post-injury, 1 month, 6 months and 18 months. PTH was assessed at visit 4 by a post-TBI headache questionnaire. Healthy controls (n=21) were scanned twice 6 months apart.

**Results:** Compared to non-PTH, PTH patients had decreased GMV across two large clusters described as the right anterior-parietal (p=0.012) and left temporal-opercular (p=0.027). Compared to healthy controls non-PTH patients had decreased GMV in the left thalamus (p=0.047); PTH patients had decreased GMV in several extensive clusters: left temporal-opercular (p=0.003), temporal-parietal (p=0.041), superior frontal gyrus (p=0.008) and right middle frontal/superior frontal gyrus (0.004) and anterior-parietal (p=0.003).

**Conclusion:** Differences between PTH and non-PTH patients were most striking at early time points. These early changes may be associated with an increased risk of PTH. Patients with these changes should be monitored for chronic PTH.

## Introduction

Mild traumatic brain injuries (mTBI) account for 75% of all brain injuries sustained ranging from 131-640 per 100,000 population, although many go unreported (Obermann et al. 2010; Obermann et al. 2009). A subset of mTBI patients develop persistent, sometimes disabling disorders, with one of the most common being post-traumatic headache (PTH) (Anderson et al. 2015; Defrin 2014; Lucas et al. 2012). PTH usually arises within seven days of the injury and is considered chronic if it persists three months after injury (Defrin 2014; Lucas et al. 2012). The prevalence of PTH has been reported as high as 79% at 3 months and 65% at 12 months following mTBI (Luca et al. 2012).

Several MRI studies have characterized the structural and functional changes associated with mTBI and its symptoms (Eierud et al. 2014). Wide spread white and grey matter reductions have been reported. In particular decreases in GMV have been reported in the thalamus, right precentral and postcentral gyri and supplementary motor area (Bendlin et al. 2008). Other studies have found that the GMV atrophy one year after TBI was located in the right precuneus, suggesting vulnerabilities in particular areas of the brain (Zhou et al. 2013).

Despite the common occurrence of PTH, its pathogenesis is unknown. The spectrum of contributing factors both biological and behavioral are likely to vary by person (Lucas et al. 2012). In chronic PTH damage to the pain modulatory system after mTBI has been implicated in the development of the headache disorder (Defrin 2014). Obermann et al. (2009) reported that PTH in patients post-whiplash was associated with significantly decreased GMV in the anterior cingulate cortex and dorsolateral prefrontal cortex (DLPFC) at two weeks and three months post-injury; however, these changes completely resolved along with the headache after a year. Studies have reported mixed results on resolution of PTH in various TBI populations and have yet to define a core set of predictors or pathophysiological changes associated with PTH (Obermann et al. 2009). Given the difficulty in assessing these predictors and changes associated with PTH in TBI, we assessed patients both with and without headache and explored GMV changes that are unique to mTBI and those specific to patients with headache post-mTBI. In this secondary analysis we aimed to compare changes in GMV in mTBI patients with and without PTH and healthy controls over four time points occurring up to 18 months after injury.

## Materials and Methods

### Participants and Study Design

Fifty-one mTBI patients (15 females, 36 males) seen at the R. Adam Cowley Shock Trauma Center at the University of Maryland Medical Center were prospectively recruited and enrolled between March 2010 and May 2012. These patients were a subset of a larger imaging protocol using a combination of advanced MR imaging and neuropsychological assessments including the Automated Neuropsychological Assessment Metrics (ANAM) and the Modified Rivermead Post-Concussion Symptom Questionnaire (RPQ) which was used to assess the level of post-concussive symptoms (Kane et al. 2007; King et al. 1995). Methods have been described in detail previously (Sours et al. 2015). In brief, patients were screened and excluded if they had a history of the following: neurological and psychiatric illness, stroke, brain tumors or seizures and contraindications to MR. All patients were classified as having a mTBI based on their Glasgow Coma Scale (GCS) score (range 13-15) and mechanism of injury consistent with trauma. Additionally, 21 healthy controls, age 18 or older, free of mTBI and no previous head injury resulting in hospitalization, were enrolled. Exclusion criteria for patients also applied to controls.

mTBI patients were seen at four time points starting approximately within 10 days post-injury, one (1) month, six (6) months, and eighteen (18) months post-TBI. For this secondary analysis of patient data, mTBI patients were classified as post-traumatic headache (PTH) based on self-reported headache post-TBI using an in-house developed post-TBI headache questionnaire. The questionnaire was administered at the final visit (approx. 18 months post injury) and was modified from the Theeler et al. (2010) headache questionnaire which assessed PTH in post-concussive soldiers. The questionnaire included, but was not limited to, the start date of headache post-TBI (within one week or within one month post-injury), headache pain intensity, quality of the headache pain (throbbing or not) and location of the headache pain (one sided or not). Controls were seen twice approximately six months apart. Each visit included an MRI scan. For the purposes of this study, only structural MRI data was utilized. This study was approved by the Internal Review Board at the University of Maryland and all participants provided written informed consent and HIPAA compliance.

### MRI Data Acquisition

Imaging was performed on a Siemens Tim-Trio 3T MRI scanner using a 12 channel receiver head coil. A high resolution image T1-MPRAGE was acquired in either axial (TE=3.44 ms, TR=2250 ms, flip angle=9°, resolution=256 × 256 × 96, FOV=22 cm, slice thickness=1.5 mm) or sagittal (TE= 2.91 ms, TR=2300 ms, flip angle 9°, resolution 256 × 256 × 176, FOV=256mm, slice thickness=1 mm, with parallel imaging factor of 2). Controls received scans with the axial protocol, but some patients had scans with the sagittal sequence (See below for details).

## Data Analysis

### Voxel Based Morphometry (VBM)

We used longitudinal VBM to compare whole-brain GMV between mTBI patients with and without PTH and healthy controls. After realignment all T1 images were preprocessed using the Computational Anatomy Toolbox (CAT12) (http://www.neuro.uni-jena.de/cat12/) in SPM12. The preprocessing pipeline consists of several steps in which the images are spatially normalized to MNI space (resampled to a voxel size of 1.5mmX1.5mmX1.5mm), segmented into grey matter (GM), white matter (WM) and cerebrospinal fluid (CSF), and smoothed with an 8mm Gaussian Kernel. Whole-brain GMV changes were calculated in both the TBI patients and the healthy controls, and an absolute threshold mask of 0.1 was employed for analyses. This threshold excludes voxels with less than 10% probability of being grey matter occurring because of partial volume and smoothing effects.

Patient scans were acquired in both axial and sagittal sequences over time; the sagittal sequence which was conducted on the same scanner, was introduced later in the study during data acquisition as it produced better quality scans. Initial analyses attempted inclusion of all patient scans regardless of acquisition sequence, but preprocessing failed. Therefore for patients with scans acquired in both the axial and sagittal sequences at different visits, had their visits with the most consistent scan sequence included in the data analysis used most often was used. E.g., if a patient was seen at all four time points and had 3 axial scans and one sagittal, we excluded the sagittal scan. Patients with equal numbers of axial and sagittal scans had their sagittal scans utilized in the analysis. This resulted in the exclusion of 25 scans from 19 patients across all four time points in the non-PTH patients and 10 scans from 8 patients in the PTH group. Given that the missingness was at random (the timing of the upgrade and the scans impacted was not systematic) and that the excluded scans spanned across all visits and both groups (affecting equal numbers in both PTH and non-PTH), the resulting sample of images was likely representative of the entire sample.

Quality control of all scans was conducted before and after preprocessing. This included visual inspection and assessing the quality of pre-processing using the CheckReg function in SPM12. Post preprocessing each image was assessed for quality using quality control tools in CAT12. Though there is no clear cut off for a bad image we decided that images which were below more than two standard deviations were further assessed in CheckReg. Due to abnormal brain morphology one participant from the PTH group was excluded from analysis, resulting in a final sample size of 50 mTBI patients.

### Sandwich Estimator Analysis (SwE)

Since all patients were not seen at every time point and some scans had to be dropped due to inconsistencies in scan type, SwE was used to account for the unbalanced nature of the data (Guillaume et al. 2014). It uses an unstructured covariance and accounts for all the random effects possible in the model and has been shown to be particularly robust for longitudinal neuroimaging data (Guillaume et al. 2014). Using the non-parametric SwE model with 999 bootstraps, group-by-time interactions were assessed (PTH vs non-PTH x visit) to examine change in GMV over time in mTBI associated with PTH and (PTH and non-PTH vs Control x visit) to examine change in GMV over time associated with mTBI and PTH. Models in SwE to compare PTH and non-PTH were adjusted for sex to account for unbalance over time due to loss to follow up. A cluster forming threshold of p=0.005 was applied and FWE (estimated from the wild bootstrap distribution) was set to 0.05 (Guillaume et al. 2014).

### Cluster and Volume Extraction for Brain Regions

Results are written in a -log10 cluster-wise and FWE-corrected non-parametric p value image. To obtain the clusters with a corrected p-value ≤0.05, the images were thresholded using the following approach in FSL cluster tool (https://fsl.fmrib.ox.ac.uk/fsl/fslwiki/Cluster): - log10(0.05)=1.301. Whereas traditional MRI clusters results are given with coordinates to identify a peak voxel, results from SwE cluster analyses are written to produce clusters where all voxels within any given cluster have the same intensity value. Each cluster has one peak value which indicates where the intensity value is first located. For visualization of the results, clusters were displayed on a brain surface using Surflce (https://www.nitrc.org/projects/surfice/).

Marsbar was used (https://www.nitrc.org/projects/marsbar/) to extract data for significant clusters identified in the above analyses. Values were converted to mm^3^ using the following equation: cluster size x voxel size x beta value. The beta value extracted represents the proportion of the voxel attributed to GM. Differences between patient groups at each time point and patient groups and controls were assessed using 2-sample T-tests and Wilcoxon tests. Mean volume over time by group was plotted and the coefficient of the interaction between group and time was analyzed using linear mixed models. Whole brain differences in GM and WM were also assessed. Using the total intracranial volume (TIV) estimation module in CAT12 the volumes for GM, WM and CSF and TIV were extracted. All statistical analyses including plots were conducted in SAS (v.9.4, SAS Institute Inc. Cary, NC). Testing was two-sided and done at the 0.05 level of significance.

### Comparison of GMV and Demographic Variables by PTH and Control Status

Linear and logistic regression analyses provided comparisons of clinical and demographic variables between mTBI patient groups and controls. Additionally differences in overall RPQ as well as anxiety and depression subscales of the RPQ were examined between the PTH and non-PTH groups. We also assessed the relationship between RPQ total scores and subscale scores with GMV, in all significant clusters in the group-by-time interaction SwE analysis. This analysis was conducted using linear mixed models in SAS (v.9.4, SAS Institute Inc. Cary, NC). Testing was two-sided and done at the 0.05 level of significance.

## Results

### Sample Description

Thirty-one non-PTH and 19 PTH patients were included in the final analysis. PTH and non-PTH mTBI patients were of similar age (p=0.41). Controls were on average 40 years old and were seen at two time points 205±42 days apart (Table 1). Total Rivermead scores were higher in PTH than non-PTH at both visits 1 (p=0.032) and 4 (p=0.002) (Table 1). In longitudinal models there was no significant relationship between GMV and overall RPQ scores or anxiety and depression subscales. Eighteen mTBI patients had evidence of head injury with positive CT results (12 non-PTH; 6 PTH). Eighteen PTH patients completed the headache questionnaire with an average headache pain of 5 on a 0-10 scale. Sixteen reported headaches within one week of injury, with all but 2 reporting headaches at the end of follow-up.

**Table 1:** Demographic and clinical characteristics of healthy controls and mTBI patients by posttraumatic headache status

| **Variable** | **Non-PTH** | **PTH** | **Healthy Controls** |
| --- | --- | --- | --- |
| Age/years | 42±16 | 48±21 | 40±19 |
| Male (n) | 24 | 11 | 11 |
| Female (n) | 7 | 8 | 10 |
| **Clinical Characteristics in mTBI patients** |  |  |  |
| **Time** | **Non-PTH** | **PTH** | **p-value\*** |
| **Days post-TBI (n) when MRI scan occurred** |  |  |  |
| Visit 1 | 6.0±3(23) | 7±3(17) | 0.27 |
| Visit 2 | 37±10(22) | 34±7(18) | 0.40 |
| Visit 3 | 193±26(29) | 206±29(17) | 0.18 |
| Visit 4 | 600±92(25) | 538±72(14) | **0.037** |
| **Total Rivermead Score** |  |  |  |
| Visit 1 | 12±16 | 21±17 | **0.032** |
| Visit 2 | 15±15 | 17±15 | 0.61 |
| Visit 3 | 9±12 | 18±20 | 0.16 |
| Visit 4 | 8±10 | 29±22 | **0.002** |

\*p values based on T-tests, Chi-Square and Wilcoxon tests. Values are Mean±S.D mTBI mild traumatic brain injury, PTH post traumatic headache

### PTH vs. Non-PTH

Total GMV over time differed between PTH and non-PTH, with PTH having an overall decrease over time (Please refer to supplementary figure 1 in the online resource). SwE analysis revealed no areas of increased GMV in PTH patients over time in the interaction model, but there was a significant group by time interaction between the two mTBI groups resulting in two extensive clusters of decreased GMV (Table 2). Figure 1a shows the region described as the left temporal-opercular cluster (p=0.027) (1905 voxels), which includes left middle temporal gyrus (MTG), superior temporal gyrus (STG) and parietal operculum, and the second cluster labeled the right anterior-parietal cluster (p=0.012). This latter cluster was even more extensive (3166 voxels) spanning areas of frontal, temporal and parietal lobes including right inferior temporal gyrus (ITG), middle temporal gyrus (MTG), angular gyrus (Ang), supramarginal gyrus, superior temporal gyrus (STG), primary somatosensory cortex (S1) and primary motor cortex (M1). Table 3 shows the difference in GMV between PTH and non-PTH at each of the four time points and figure 1a also shows plots of the change in GMV in these regions over time in PTH and non-PTH. In the left temporal-opercular cluster the difference between the two groups is significant only at visit 1 (p=0.022), while in the right anterior-parietal cluster differences were most evident at visits 2 and 4. However, PTH consistently show negative slopes over time, starting with higher GM volumes compared to non-PTH and decreasing through to visit 4 in both areas. This change in GMV was significant in PTH in both clusters (p<0.0001) as indicated by the interaction p-value whereas in non-PTH the change over time was relatively constant and not significant.

**Table 2:**
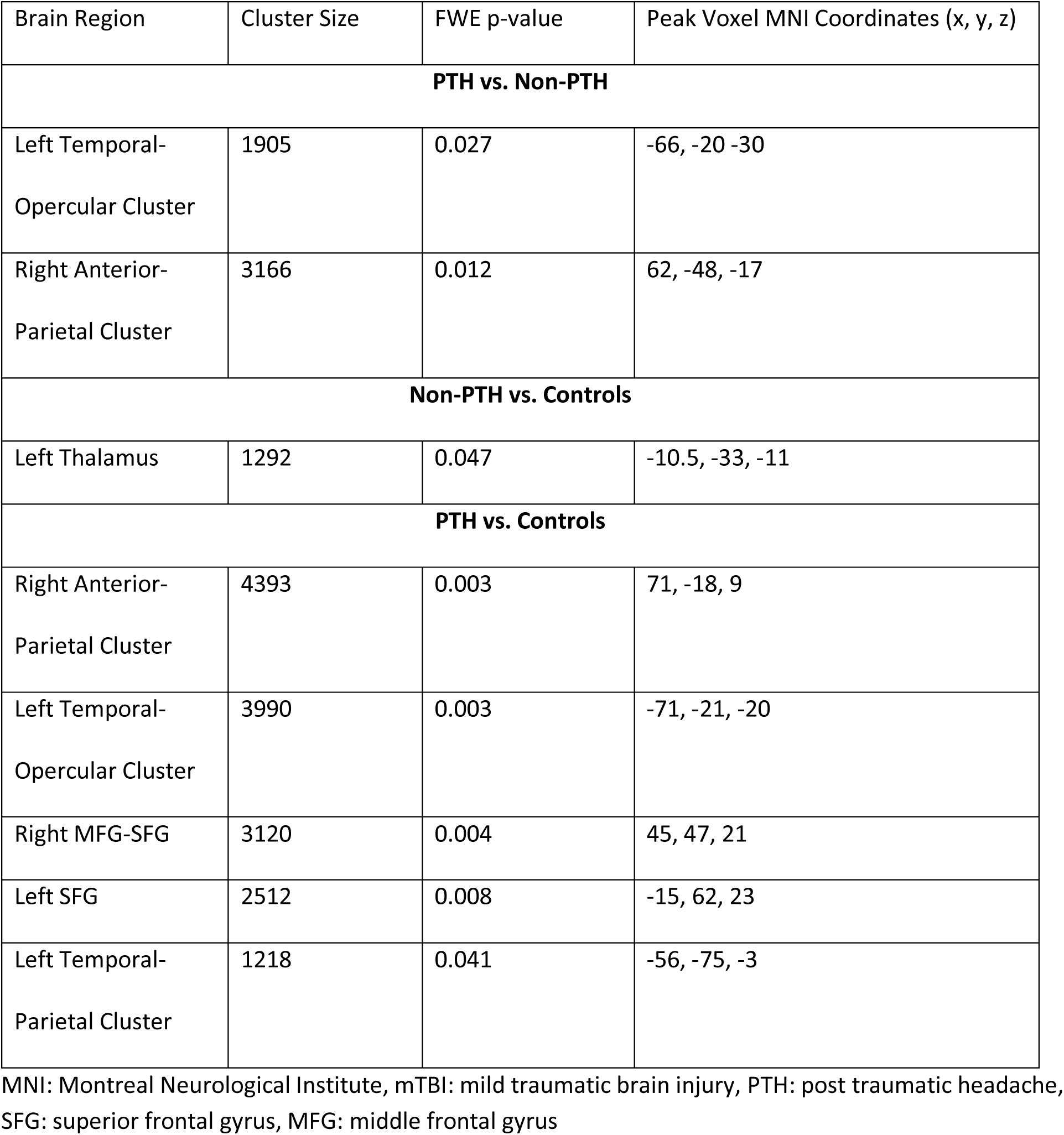
Significant clusters from whole brain time by group analysis.

**Fig. 1.**
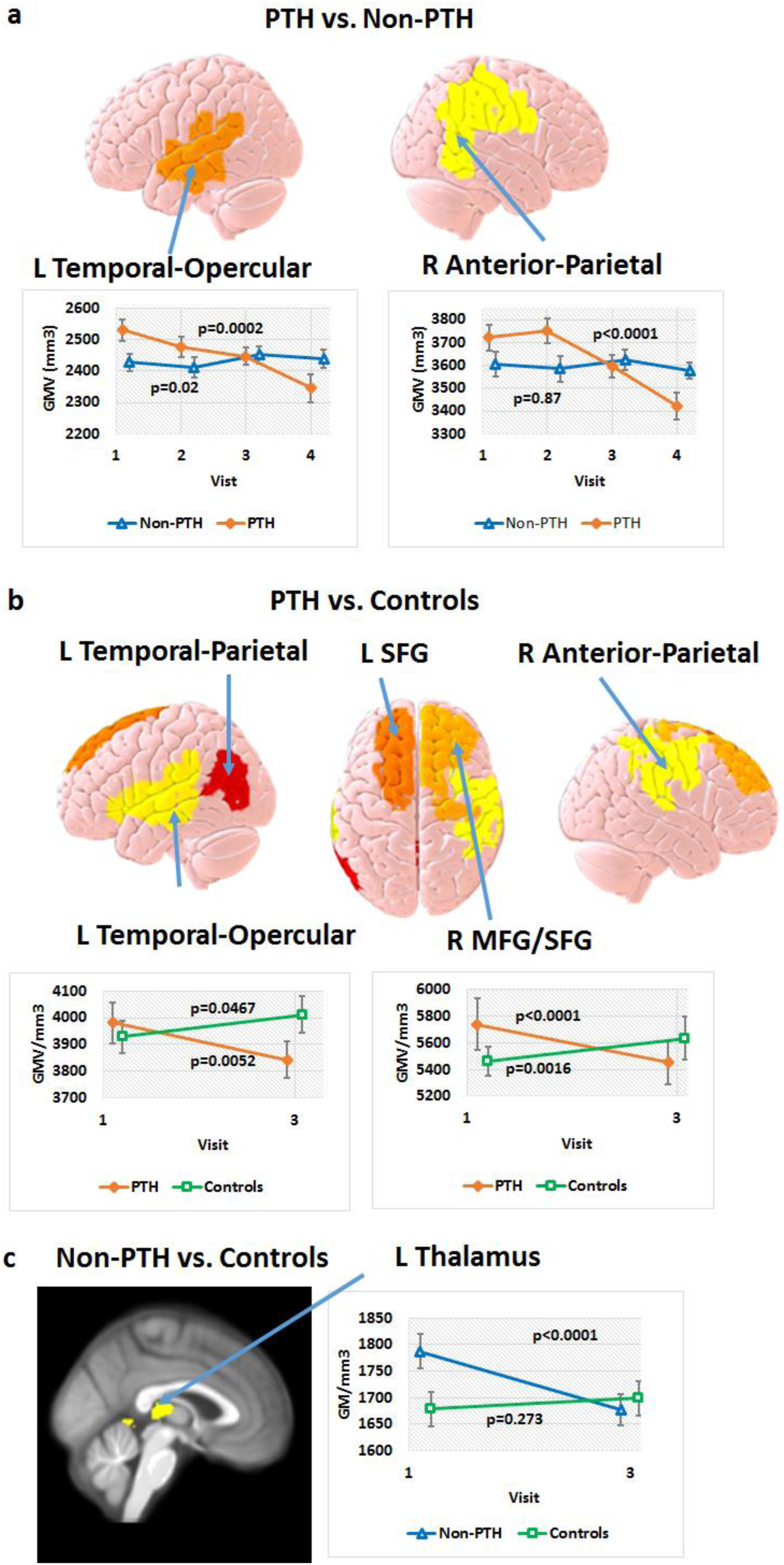
Decreased GMV in mTBI patients over time. Results based on SwE analyses. **A)** decreased GMV over time in mTBI patients with PTH compared to non-PTH in the left temporal-parietal and right anterior parietal regions. **B)** decreased GMV over time in mTBI patients with PTH compared to healthy controls in the left temporal-opercular, temporal-parietal cluster, superior frontal gyrus (SFG) and right middle/superior frontal gyrus (MFG/SFG) and anterior parietal. Plots are shown for the Temporal-Opercular and MFG/SFG clusters. **C)** decreased GMV over time in mTBI patients with non-PTH compared to healthy controls in the bilateral thalamus. A cluster forming threshold of p<0.005 was applied and FWE (estimated from the wild bootstrap distribution) was set to 0.05. In all plots PTH patients are represented by an orange line with filled diamond markers. Non-PTH are represented by a blue line with triangle markers and controls are represented by a green line with square markers. Error bars in plots represent the standard error and are centered at the mean for each group at each time point. P values represent the interaction term for group*time from a linear mixed model.

### mTBI vs. Healthy Controls

PTH had decreased GMV compared to healthy controls in five clusters (Figure 1b, Table 3), two of which were similar to those observed between PTH vs. non-PTH. These clusters included left temporal-opercular cluster (p=0.003), temporal-parietal cluster (p=0.041) and the SFG (p=0.008). The left temporal-opercular cluster covered the parietal and frontal operculum, STG, and MTG. The temporal-parietal cluster included the angular gyrus and the parietal junction. There was also decreased volume in two clusters on the right: a middle/superior frontal gyrus cluster (MFG/SFG) (p=0.004) and an anterior-parietal cluster (p=0.003). The MFG/SFG cluster extends posteriorly to M1. The right anterior-parietal cluster includes the STG, supramarginal gyrus, M1 and S1 (Figure 1b).

**Table 3:** Grey matter volumes extracted from significant clusters in mTBI patients by posttraumatic headache status

| Time | Non-PTH | PTH | Difference | p value* |
| --- | --- | --- | --- | --- |
| **Left Temporal-Opercular Cluster** |  |  |  |  |
| Visit 1 | 2426.7± 128.6 | 2529.9± 143.3 | -103.2 | **0.022** |
| Visit 2 | 2411.1± 156.3 | 2476.1± 139.2 | -64.9 | 0.18 |
| Visit 3 | 2451.1± 146.0 | 2447.0± 112.0 | 4.1 | 0.92 |
| Visit 4 | 2439.9± 145.0 | 2344.8± 164.9 | 95.1 | 0.069 |
| **Right Anterior-Parietal Cluster** |  |  |  |  |
| Visit 1 | 3605.6± 255.1 | 3721.9± 234.9 | -116.2 | 0.15 |
| Visit 2 | 3584.9± 258.8 | 3752.4± 230.4 | -167.5 | **0.039** |
| Visit 3 | 3623.9± 253.0 | 3594.8± 204.7 | 29.1 | 0.69 |
| Visit 4 | 3575.3± 180.8 | 3419.7± 221.0 | 155.7 | **0.022** |

\*p values obtained from post-hoc T-test. Grey matter volumes (mm<sup>3</sup>) based on cluster result from whole brain time by group analysis from SwE. Values are Mean±S.D. mTBI mild traumatic brain injury, PTH post traumatic headache

Non-PTH had decreased GMV compared to controls in one cluster, which covered the bilateral thalamus and extended to the left and right parahippocampal gyrus (p=0.047; Table 3, Figure 1c). The decrease over time was also noted in PTH, but it was not significantly different from controls as PTH started with lower thalamus volume than non-PTH (Please refer to supplementary figure 2 in the online resource).

Figures 1b and 1c show GMV change over time in each mTBI group compared to controls. PTH show a consistently negative slope in all regions. Thalamic GMV was consistent in controls over time (p=0.27) but is initially higher in the non-PTH group and decreases significantly over time (p<0.0001). PTH and controls start with comparable GMV in the left temporal-parietal and SFG clusters, while controls increase over time, PTH patients decrease between visits 1 and 3 (p<0.0001) (plots not shown). In the left temporal-opercular cluster, right anterior-parietal, and MFG/SFG clusters, PTH initially had larger volumes which decrease over time compared to controls (who have an increase in volume). Though differences between PTH patients and controls were not significant except for the right anterior-parietal cluster at visit 3 (Please refer to supplementary table 1 in the online resource), the change within groups over time was significant. mTBI patients regardless of headache status did not show increased GMV compared to healthy controls.

## Discussion

We found that mTBI patients with evidence of PTH approximately 18 months post-injury had decreased GMV in two main regions of the brain compared to mTBI patients who do not develop headache. These were defined as the left temporal-opercular cluster and the right anterior-parietal cluster, and in both there was a steady GMV decline over time in PTH, but relatively stable GMV in the non-PTH group. Differences between the two groups were apparent at visit 4 in the left temporal-opercular cluster and at visit two in the right anterior-parietal cluster.

Compared to controls, both mTBI groups showed a general decrease in GMV over time. Controls were only scanned at two time points, so comparisons to patients could only be made between visits 1 and 3. Therefore the observed decrease in PTH patients in the R MFG/SFG may be attributed to patients having initially higher GMV at visit 1 which declined to comparable levels to controls at visit 3. Conversely in the left temporal-opercular cluster, PTH patients and controls have comparable volumes at visit 1 and diverge by visit 3. The anterior-parietal cluster, which is decreased in PTH compared to both non-PTH and controls, extends into the somatosensory cortex, which is often implicated in headache disorders (DaSilva et al. 2007).

Non-PTH patients compared to controls had decreased thalamic GMV over time. PTH patients showed a similar, non-significant trend. Thalamic GMV loss post-TBI up to one year after injury has been reported, and this trajectory differs from healthy controls (Eierud et al. 2014; Lucas et al. 2014), consistent with our findings in non-PTH patients.

Research shows that persistent PTH patients compared to healthy controls had decreased cortical thickness in several regions including bilateral SFG, caudal MFG, and precentral gyrus, right supramarginal, superior and inferior parietal and precuneus regions (Chong et al. 2018). Persistent PTH was also associated with decreased cortical thickness compared to migraine patients and healthy controls in the right orbitofrontal cortex, right supramarginal gyrus and left superior frontal lobe (Schwedt et al. 2017). These are consistent with our finding where PTH had decreased GMV over an extensive area, including MFG and bilateral SFG, compared to controls and in the right anterior-parietal cluster, including supramarginal gyrus, compared to controls and non-PTH. Furthermore the decreased GMV in our PTH patients in the parietal region is consistent with the cortical thickness results found in those with persistent PTH (Chong et al. 2018). While few studies on GMV changes exist for PTH, there are many on migraine and other headache disorders. A recent meta-analysis in migraine reported decreased GMV in bilateral inferior frontal gyrus (IFG), left MFG, cingulate gyrus and right precentral gyrus (Jia and Yu 2017). PTH and migraine have similar phenotypes even though PTH is a secondary headache disorder (Schwedt et al. 2017). Results from the current study and others suggest that GMV changes in some brain areas might be specific to PTH and not migraine or headache disorders more generally, but further work is required to clarify this.

A strength of our study compared to recent research is the longitudinal follow-up of our patients. Not only do we report decreased GMV over time when compared to healthy controls and non-PTH patients, but our results show that these differences are present in PTH patients as early as one week post-TBI. Though the pathology of PTH is still unclear research suggests it may involve peripheral and central mechanisms (Defrin 2014). In mice following a mild closed injury, enhanced peripheral cranial nociception is observed which may be associated with the initiation of PTH (Benromano et al. 2015). The initial increased GMV that we observed, particularly in regions of the somatosensory cortex in PTH patents, may represent damage to the pain processing areas or pathways, increasing the vulnerability to the development of PTH. Conversely, decreased GMV over time is consistent with an effect of ongoing pain experienced by patients in our sample.

While our study provides some important insights into PTH, there were some limitations. The sample consisted of 19 PTH patients, some of whom were not seen at all four time points. However, by comparing PTH and non-PTH mTBI patients as well as healthy controls we were able to assess both the impact of mTBI and PTH on GMV. The use of the SwE toolbox allowed us to account for the unbalanced nature of the sample across visits and perform non-parametric inference. Furthermore, the original study was not developed to study PTH, so several methodological considerations that would have improved the conclusions drawn were not implemented. By assessing headache at 18 months via a patient-completed questionnaire there was possible misclassification of persons by headache group. We were also unable to capture patients as they transitioned from acute to no headache or to chronic headache. While this co-mingling of headache status likely biases the result towards a null finding as patients with resolved headache likely recovered GMV, it is an important nuance in the interpretation of the results. By including patients with both acute and chronic PTH in the study, we were able to capture those patients who had greater morbidity due to PTH and examine the effects on their GMV. While this may make it difficult to compare our results to other PTH studies, it highlights the fact that many mTBI patients experience PTH longer than 3 months post-injury. Future studies will therefore be aimed at recruiting larger numbers of mTBI patients and recording PTH status at multiple time points (as well as ascertain pre-existing headache status), including at one week and 3 months, as a means to capture progression of PTH and its relationship with GMV (Lucas et al 2012; Stacey et al. 2016).

The original study did not assess participant headache status prior to injury. If there were patients with pre-existing migraine or any other headache disorder prior to injury this could bias the results. Since the questionnaire to ascertain headache status was administered at the final visit, there is a possible recall and selection bias as only those seen at that visit would have received the questionnaire and those who had PTH at that visit are likely to have differential recall of headache over the study period.

The literature on the impact of mTBI and PTH on brain structure is growing, and our findings are consistent with cortical thickness studies in persistent PTH. By following patients for 18 months, we were able to capture GMV differences at the first visit post-injury and changes over time. We found brain regions in PTH with decreased GMV compared to both healthy controls and non-PTH patients. Some of these differences were evident as early as one week post-injury, suggesting that early brain changes following mTBI might predict risk for PTH.

## Supporting information

## Acknowledgements

We thank Thomas Nichols PhD and Bryan Guillaume PhD for their technical and analytical support with SwE Toolbox and Christian Gaser PhD for the development and provision of the CAT12 Toolbox. Funding: Department of Defense (W81XWH-08-1-0725 & W81XWH-12-1-0098 to RPG), NIH/NCCIH (R01 AT007176-01A1 to DAS). The authors declare no competing financial interests.

