## Supplementary material for "Decreased grey matter volume in mTBI patients with post-traumatic headache compared to headache-free mTBI patients and healthy controls: a longitudinal MRI study"

**Supplementary Table 1: Grey matter volumes for significant clusters comparing mTBI patients and healthy controls by posttraumatic headache status**

| Time | No PTH | Healthy Controls | Difference | p value* |
|---|---|---|---|---|
| **Left Thalamus** | | | | |
| Visit 1 | 1787.3±154.9 | 1678.4±147.8 | 108.9 | **0.022** |
| Visit 3 | 1676.3±157.9 | 1698.5±149.0 | -22.3 | 0.62 |
| **Time** | **PTH** | **Healthy Controls** | **Difference** | **p value*** |
| **Left Temporal-Parietal Cluster** | | | | |
| Visit 1 | 1715.0±157.2 | 1690.2±1690.2 | 24.8 | 0.61 |
| Visit 3 | 1636.2±149.7 | 1716.4±155.2 | -80.2 | 0.12 |
| **Left SFG Cluster** | | | | |
| Visit 1 | 3981.3±314.4 | 3928.6±273.6 | 52.7 | 0.58 |
| Visit 3 | 3841.8±281.2 | 4011.9±318.8 | -170.1 | 0.093 |
| **Left Temporal-Opercular Cluster** | | | | |
| Visit 1 | 6640.1± 545.0 | 6459.6± 424.8 | 180.5 | 0.26 |
| Visit 3 | 6385.5± 510.8 | 6591.0± 494.5 | -205.4 | 0.22 |
| **Right Anterior-Parietal Cluster** | | | | |
| Visit 1 | 6092.8± 496.4 | 5843.9± 325.4 | 248.8 | 0.071 |
| Visit 3 | 5766.9± 378.8 | 6131.7± 343.8 | -364.8 | **0.004** |
| **Right MFG-SFG Cluster** | | | | |
| Visit 1 | 5738.2±795.5 | 5461.9±507.2 | 276.3 | 0.20 |
| Visit 3 | 5450.1±657.7 | 5633.4±743.0 | -183.3 | 0.43 |

*p values obtained by post-hoc T-test. Grey matter volumes ($mm^3$) are based on cluster result from whole brain time by group analysis from SwE. Values are Mean±S.D. SFG superior frontal gyrus, MFG middle frontal gyrus.

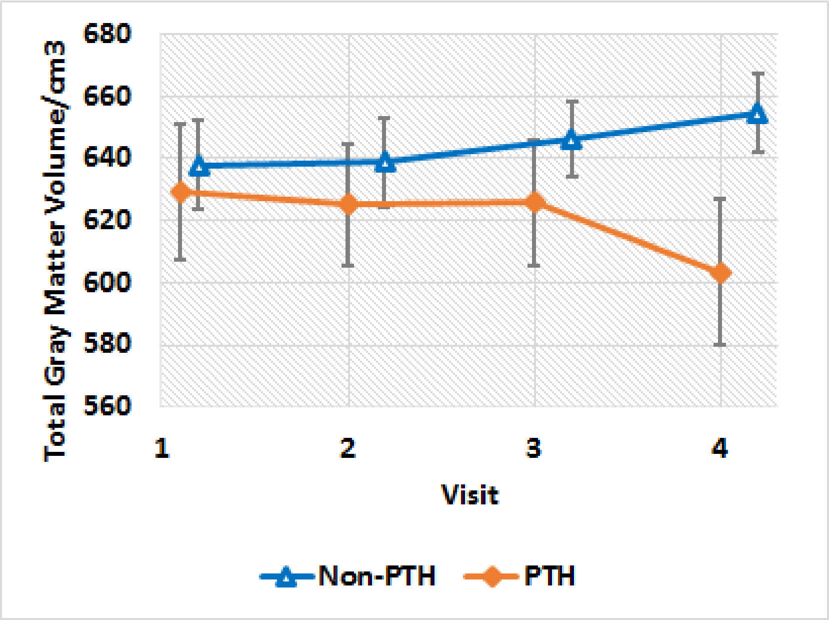

**Supplementary Fig. 1** Change in total gray matter volume over time in PTH and non-PTH patients. Error bars represent the standard error and are centered at the mean for each group at each time point. Non-PTH patients are represented by the blue line with triangle markers and PTH patients are represented by the orange line with filled diamond shaped markers.

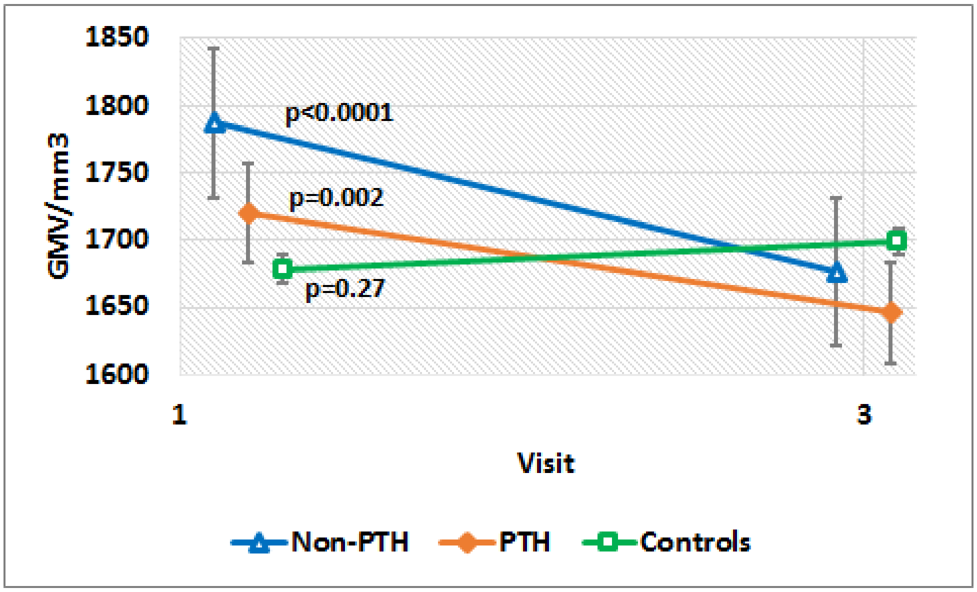

**Supplementary Fig. 2** Change in thalamic gray matter volume over time in mTBI patients (PTH and non-PTH) vs. Controls. Error bars represent the standard error and are centered at the mean for each group at each time point. A cluster forming threshold of $p<0.005$ was applied and FWE (estimated from the wild bootstrap distribution) was set to 0.05. P values represent the interaction term for group*time from a linear mixed model. Non-PTH patients are represented by the blue line with triangle markers, PTH patients are represented by the orange line with filled diamond markers and controls are represented by the green line with square unfilled markers.
